# Towards guided mutagenesis: Gaussian process regression predicts MHC class II antigen mutant binding

**DOI:** 10.1101/2021.04.14.439878

**Authors:** David R. Bell, Serena H. Chen

## Abstract

Antigen-specific immunotherapies (ASI) require successful loading and presentation of antigen peptide into the major histocompatibility complex (MHC) binding cleft. One route of ASI design is to mutate native antigens for either stronger or weaker binding interaction to MHC. Exploring all possible mutations is costly both experimentally and computationally. To reduce experimental and computational expense, here we investigate the minimal amount of prior data required to accurately predict the relative binding affinity of point mutations for peptide-MHC class II (pMHCII) binding. Using data from different residue subsets, we interpolate pMHCII mutant binding affinities by Gaussian process (GP) regression of residue volume and hydrophobicity. We apply GP regression to an experimental dataset from the Immune Epitope Database, and theoretical datasets from NetMHCIIpan and Free Energy Perturbation calculations. We find that GP regression can predict binding affinities of 9 neutral residues from a 6-residue subset with an average R^2^ coefficient of determination value of 0.62 ± 0.04 (±95% CI), average error of 0.09 ± 0.01 kcal/mol (±95% CI), and with an ROC AUC value of 0.92 for binary classification of enhanced or diminished binding affinity. Similarly, metrics increase to an R^2^ value of 0.69 ± 0.04, average error of 0.07 ± 0.01 kcal/mol, and an ROC AUC value of 0.94 for predicting 7 neutral residues from an 8-residue subset. Our work finds that prediction is most accurate for neutral residues at anchor residue sites without register shift. This work holds relevance to predicting pMHCII binding and accelerating ASI design.

## Introduction

Major histocompatibility complex class II (MHCII) antigen binding and presentation to the T-cell receptor (TCR) of CD4^+^ T-cells represents a critical immunological interaction with dysfunction being implicated in autoimmune diseases such as Type-1 Diabetes^1-3^, Celiac Disease^4-5^, and Multiple Sclerosis^6-7^. Antigen specific immunotherapies (ASI) such as vaccines and HLA blockers work by modulating this MHCII-antigen-TCR interaction^4, 8-9^. Oftentimes, a known antigen is used as a template for ASI design, which is then mutated for enhanced or reduced binding interaction with MHCII and/or TCR^10-11^. Computational techniques, such as sequence-based^12-13^, or structure-based^14-18^ methods, aim to predict native antigen-MHC binding and mutations to accelerate ASI design and limit costly mutagenesis experiments.

Gaussian Process (GP) regression is a powerful Bayesian supervised machine learning method, which innately provides uncertainty estimates of predictions^19^. GP regression has been used widely in diverse fields such as geostatistics^20^ and imaging microscopy^21^. For biological applications, GP regression has been used to predict protein stability^22-23^ and turnover^24^, protein structure prediction^25^ and optimization^26-27^, as well as protein-protein binding^22, 28^ and quantitative structure activity relationship (QSAR) models for therapeutic design^29-31^. GP regression has also been used to study MHC class I antigen binding from sequence^32-33^. These models, however, have not been extended to the arguably more challenging task of predicting MHCII antigen peptide (pMHCII) binding, with ambiguous registers and variable sized epitopes, nor of predicting the mutational landscape at specific sites. We note that many sequence-based MHC-binding prediction models make use of complex machine learning architectures^13, 34^, and that neural networks, in the limit of infinite network width, are equivalent to GP models^35^. Hence, the extension of GP models for pMHCII-binding prediction and characterization of mutational landscapes is a logical outgrowth of current affinity prediction methods.

In this work, we explore the minimal data required to predict relative binding affinities of pMHCII antigen mutants using GP regression across residue volume and hydrophobicity. We study both experimental and theoretical binding affinity datasets from the Immune Epitope Database and Analysis Resource (IEDB), NetMHCIIpan-4.0 server, and Free Energy Perturbation (FEP) calculations. We find that GP regression can accurately predict pMHCII mutant binding affinities for neutral residues at anchor residue sites. More specifically, we find that GP regression across 6- and 8-residue subsets can accurately capture the binding affinities of remaining neutral residues. The corresponding R^2^ coefficient of determination values are 0.61 ± 0.04 and 0.69 ± 0.04 (± 95%CI), respectively, with AUC values of 0.92 and 0.94 for binary classification of residues with either enhanced or diminished binding affinity. Finally, we discuss how GP regression can be used to direct mutagenesis experiments and FEP calculations for increased efficiency, which provides opportunities to accelerate MHCII ASI design.

## Results

Figure 1 presents the MHCII-antigen system together with the Gaussian process (GP) affinity prediction workflow. MHCII is a heterodimer consisting of HLA-α and HLA-β protein chains. These chains construct an antigen binding cleft from two α-helices and one β-sheet, with the antigen binding between the helices. Unlike MHCI, MHCII antigens range in length from 9 to 25 residues, with most around 15 residues^36^. A 9-residue binding core epitope of the antigen binds deeply into the MHCII binding cleft, while the flanking domains contact exterior portions of MHCII. From this 9-residue epitope, 4 anchor residues bury deeply into the MHCII (p1, p4, p6, and p9) while the remaining 5 residues point toward the TCR. For ASI design, antigen mutations, such as the Serine (Ser) to Isoleucine (Ile) mutation of one residue illustrated in Fig. 1B, are tested to identify peptide sequences with desired change in binding affinity.

**Figure 1.**
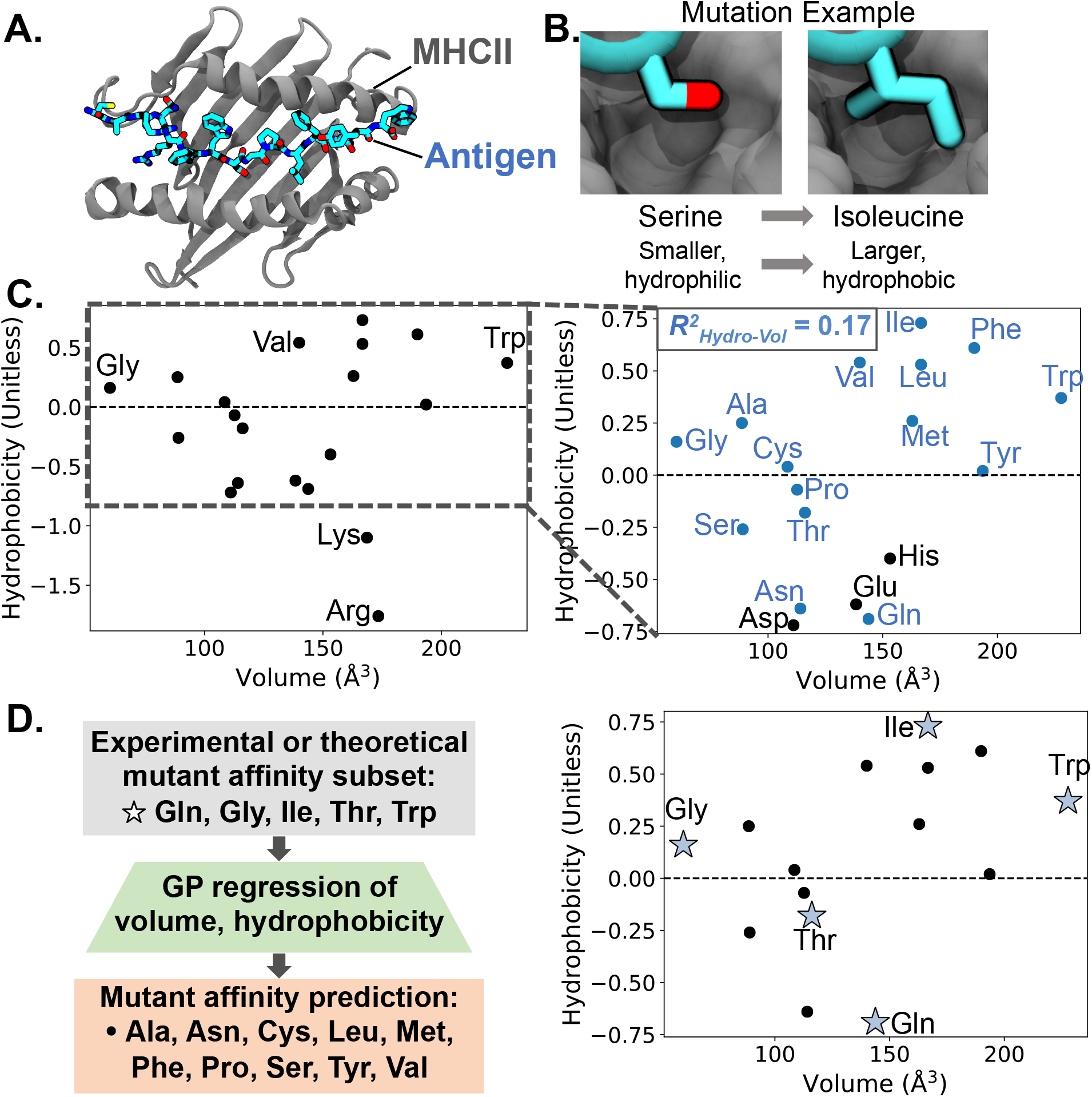
MHCII-antigen structure and Gaussian process (GP) prediction framework. **A**. MHCII in complex with antigen CARQ**<u>RFWSGPLFD</u>**YW from ref^2^ with epitope in bold underline. Only the α1 and β1 HLA domains are shown for clarity. **B**. Example antigen mutation of anchor residue Serine to Isoleucine. MHC shown in gray, antigen in cyan. **C**. Twenty standard amino acids presented by Eisenberg hydrophobicity values and residue volume in Å^3^. (Inset, right) Close-up view of amino acid plot. Pearson R^2^ correlation between hydrophobicity and volume for neutral residues (in blue) is shown in the top left corner. Pearson R^2^ correlation for all 20 standard amino acids is 0.01. **D**. GP workflow: use an experimental or theoretical mutant binding affinity residue subset (★,right) to train GP regression model across residue volume and residue hydrophobicity and then predict mutational binding affinities for the remaining residues (•, right).

For this work, we distinguish amino acids by 3 properties: residue volume measured in Å^3^ taken from ref^37^, residue hydrophobicity according to consensus Eisenberg hydrophobicity values taken from ref^38^, and residue charge. The two-dimensional landscape of residue volume and residue hydrophobicity as shown in Fig. 1C is sufficient to distinguish the 20 standard amino acids. In addition, there is low linear correlation between residue volume and residue hydrophobicity. The Pearson’s correlation R^2^ value between the two features for all 20 residues and 15 neutral residues are 0.01 and 0.17, respectively.

The GP framework we employ for this work is shown in Fig. 1D. We start with a subset of previously determined mutant residue binding affinities, either from experiment or computation, for a residue of a particular MHCII antigen. We then generate a GP model across residue volume and hydrophobicity, fitting to the known affinity values. Lastly, we use the fitted GP model to predict binding affinities of the remaining residues. For example, we start with a 5-residue subset of Gln, Gly, Ile, Thr, and Trp binding affinities for residue site 4 in a particular antigen. We next fit a GP model across residue volume and hydrophobicity to the five known binding affinities. We then use the GP model to predict the binding affinities for the remaining 15 standard residues: Ala, Arg, Asn, Asp, Cys, Glu, His, Leu, Lys, Met, Phe, Pro, Ser, Tyr, and Val. After initial testing of all 20 standard amino acids, we found that excluding charged residues improved predictive results (see Fig. S1). Charged residues are distinguishable on the 2-D plane of residue volume-hydrophobicity, but long-range electrostatic interactions between the antigen and MHCII may affect the binding site conformation and shift registers^10, 39-40^, lowering GP prediction accuracy. Hence, we excluded charged residues and only focused on the 15 standard neutral residues for GP prediction. Although our GP models are built only across two dimensions (volume and hydrophobicity), there is an implicit third dimension of charge, which we are accounting for by only targeting neutral residues.

Figure 2 presents GP regression results for Free Energy Perturbation (FEP)-predicted binding affinities. The pMHCII systems are the X-idiotype and healthy control antigens taken from Ahmed et al.^2^ with sequences shown in Fig. 2A-B. We focused on mutations of the anchor residues Tyr6 of the X-idiotype and Ser4 of the healthy control antigens, as highlighted in magenta boxes in Fig. 2A-B. We first computed all neutral residue relative binding affinities using FEP as described previously^2, 15^. Proline was excluded due to its combined sidechain-backbone structure creating an ambiguity in sidechain mutation for the FEP method. We next studied the accuracy of all possible residue combination subsets and prediction sets for GP regression models. For instance, for a subset size of k=5 residues, we tested a total of *C(n=14,r=5) =* 2002 combinations, including a subset of Ala, Cys, Phe, Gly, Ile residues to predict Leu, Met, Asn, Gln, Ser, Thr, Val, Trp, Tyr residue affinities, a subset of Cys, Phe, Gly, Ile, Leu residues to predict Ala, Met, Asn, Gln, Ser, Thr, Val, Trp, Tyr residues affinities, etc. We then looked at the distribution of GP model accuracy compared to FEP-predicted affinities as measured by R^2^ coefficient of determination values, and error, as shown in Fig. 2A-B. We found that R^2^ of the highest scoring models reached 0.82 and 0.71 with error below 1 kcal/mol by subset size k=6 residues for both antigen systems. An important caveat of the FEP data is that from previous studies^41^, FEP-predicted affinities only agree to within 1 kcal/mol of experimental values, so GP-predicted values to within 1 kcal/mol error agrees to the highest-accuracy of the FEP method. To further illustrate the highest performing GP models, we present the GP-interpolated free energy landscapes from k=6 residues in Fig. 2C-D as well as the GP landscapes using the complete set of 14 neutral residues (absent Proline) in Fig. 2E-F. Although the complete landscapes hold more detail, the GP-interpolated landscapes from residue subset data (Fig. 2C-D) largely capture overall free energy trends.

**Figure 2.**
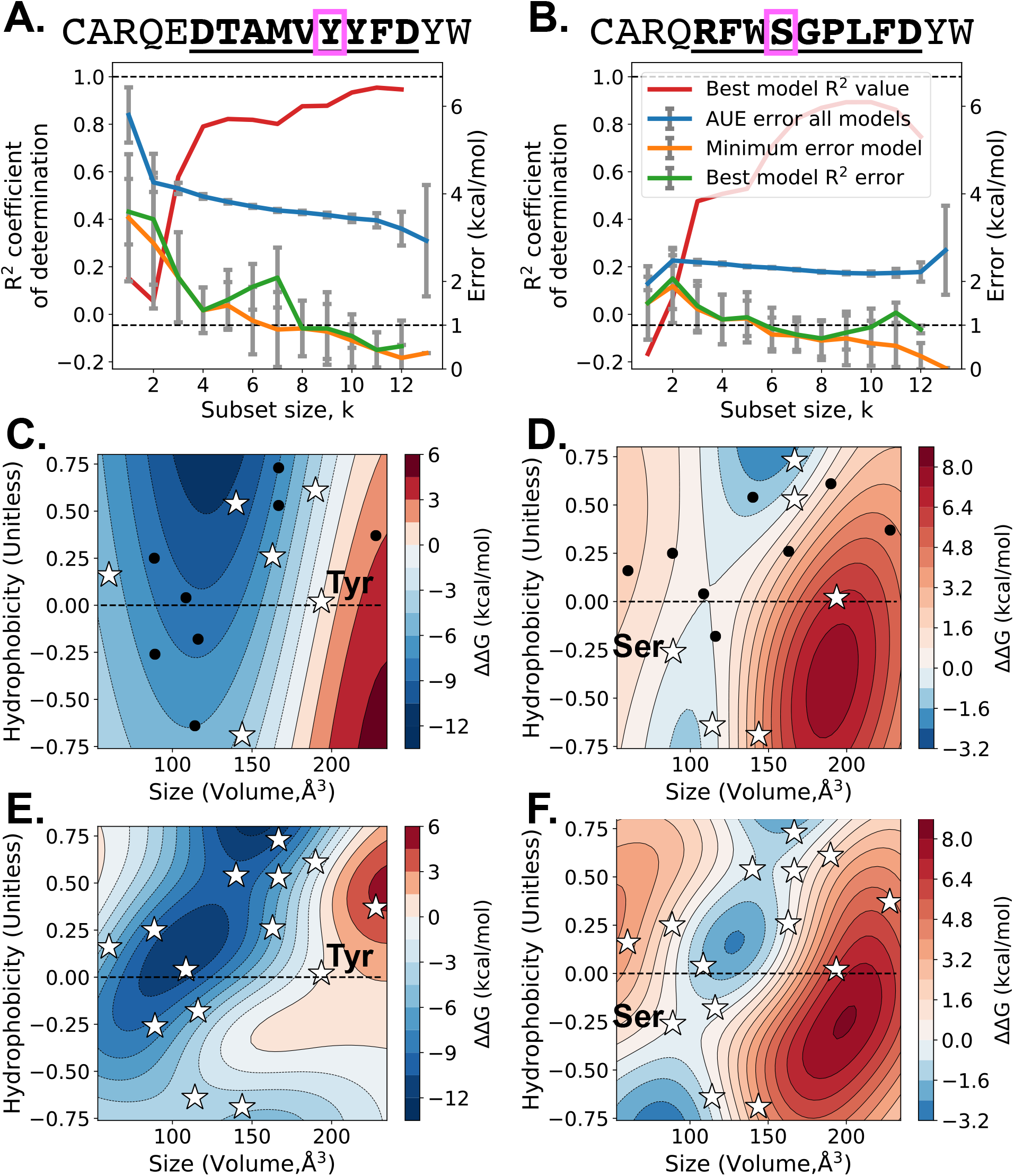
Gaussian process regression for Free Energy Perturbation-calculated binding affinities. **A-B**. Gaussian process regression of mutant binding affinities for residues boxed in magenta. Epitope sequences are in bold underline and are taken from ref^2^. (Plots) Gaussian process regression error and R^2^ coefficient of determination for all neutral residues, excluding Proline. For k=13 subset size, only 1 residue ΔG value is predicted, leaving R^2^ undefined. Error bars are 95% CI. **C-D**. Interpolated Gaussian process free energy surfaces of neutral residues constructed from a 6-residue subset (shown as white stars, with native residue labeled). Surfaces represent maximum R^2^ models. **E-F**. Gaussian process free energy surfaces of neutral residues constructed from all neutral residues (shown as stars, excluding Pro). **A,C,E**. Tyr6 mutations of CARQE**<u>DTAMVYYFD</u>**YW. **B,D,F**. Ser8 mutations of CARQ**<u>RFWSGPLFD</u>**YW.

Figure 3 presents GP prediction results using mutant affinity data from the Immune Epitope Database and Analysis Resource (IEDB)^42^. For details on the IEDB dataset, see Methods. From Fig. 3A, the overall R^2^ coefficient of determination values are lower than the FEP dataset; however, the prediction error remains low. Also seen in Fig. 3A is the large difference between the maximum R^2^ models and the average of the highest scoring models, indicating that high accuracy can be achieved, but is limited over the average. We explored this difference in prediction accuracies by generating scatterplots of the highest scoring models at k=12, as presented in Fig. S2. We found that neither the inner-quartile range of affinity values, ΔG IQR, nor the residue position show a marked trend with high scoring R^2^ models.

**Figure 3.**
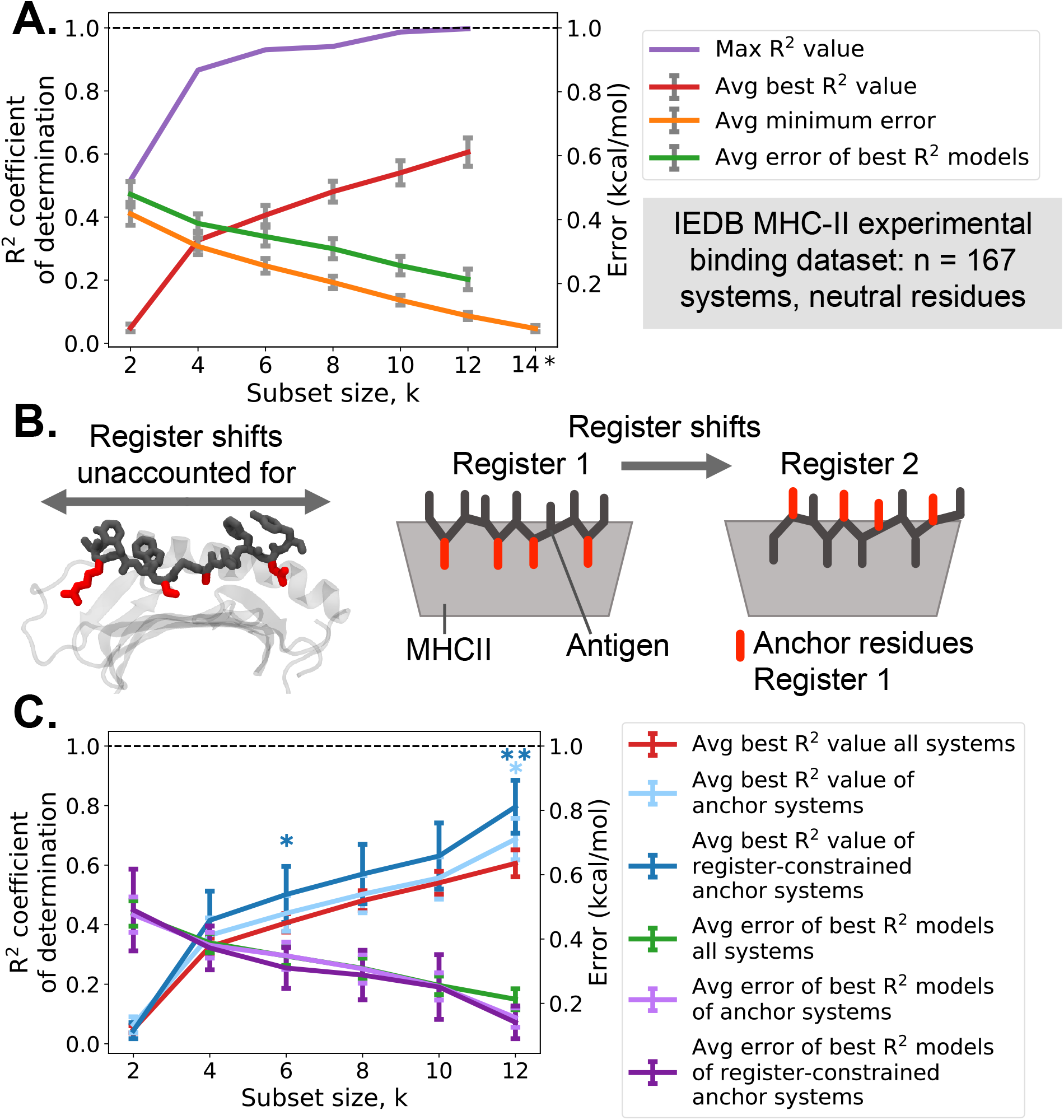
Gaussian process regression for experimentally determined binding affinities from the Immune Epitope Database (IEDB). **A**. Gaussian process regression R^2^ coefficient of determination and error for 15 neutral residues. *k* is the subset size used for prediction, e.g., for k=4, 4 residue ΔG values were used to predict the remaining 11 residue ΔG values. Averages are taken across the n=167 systems. *Subset size k=14 is excluded as only 1 residue ΔG value is predicted, leaving R^2^ undefined. **B**. IEDB binding affinity values do not account for register shifts, where the antigen moves in the MHCII binding cleft, resulting in a different binding conformation and different anchor residues (red) binding to MHCII. **C**. Gaussian process regression R^2^ coefficient of determination and error for all systems and select subsets of anchor residues and register-constrained anchor residues. * and ** indicate statistical significance with p<0.05 and p<0.005 for a one-sided t-test with null hypothesis: the average best R^2^ value of the anchor (or register-constrained anchor) set is not greater than the average best R^2^ of all sets. Accounting for register shifts as well as focusing on the anchor residues improved the accuracy of the models. All error bars are 95% CI.

We suspected that structural properties inherent in the pMHCII system but not accounted for in the IEDB dataset might explain the difference in high- and low-accuracy models. For instance, though MHCII antigens are ∼15 residues long, the 9-residue core epitope, termed the antigen register, has the greatest contact with the MHCII core binding cleft. Furthermore, in this 9-residue core epitope, residue positions 1, 4, 6, and 9, termed anchor residues, maintain large interactions with MHCII, because these residues sidechains are buried into deep MHCII binding pockets. Importantly, the IEDB does not provide information on which register the antigen is binding, nor which residues are anchor residues. If the register is shifted by one position, the anchor residues as well as the TCR-facing residues will completely change, as shown in Fig. 3B. For the GP regression we calculate here, we implicitly assume that the binding register will not change upon mutation: mutations capture the same MHCII residue environment, and the remainder of the binding surface remains unchanged. The IEDB data is not register constrained, therefore the mutations we are interpolating may, in fact, be entirely different binding surfaces. To investigate further into this hypothesis, we compared average GP model results for all systems to GP model results from anchor residues and register-constrained anchor residues, as shown in Fig. 3C. We used the NetMHCIIpan 4.0^12, 34^ webserver to predict binding registers for all 167 IEDB systems, and then selected wild-type anchor residue systems (n=42) based on residue position and register-constrained anchor residue systems (n=13) based on systems where the binding register did not change for all 15 neutral mutations. We found that GP models of register-constrained anchor residue systems were the most accurate, followed in accuracy by anchor residue systems, and lastly all systems combined. Given the favorable results of building GP models for anchor and register-constrained systems, we next sought to further explore GP model accuracy for these systems using a larger NetMHCIIpan 4.0 dataset.

In addition to FEP-calculated and experimentally determined binding affinity datasets, we further compared the prediction accuracy of GP regression with sequence-based computing methods. Figure 4 presents results from GP regression prediction of register-constrained binding affinities predicted by the NetMHCIIpan 4.0 webserver. In this analysis, we assume NetMHCIIpan predictions to be ‘ground truth’ values in order to compare the two methods; however, we note that NetMHCIIpan is itself a predictive method. To exclude the effect of possible register shifts on prediction results, for the NetMHCIIpan dataset, we only predicted mutant binding affinities of the 9-residue core epitope without the flanking domains, thereby implicitly constraining the binding register. Further, to ensure we are capturing the binding interaction between the MHCII and the antigen, we limited our dataset to anchor residue positions. The antigens selected for NetMHC prediction included antigen/MHCII systems from the IEDB dataset as well as systems implicated in Type-1 Diabetes^2, 43^. For more details on the NetMHCIIpan dataset, see Methods. Following our IEDB analysis, we focused on neutral mutations of anchor residues. We found that average best R^2^ coefficient of determination values were greater than 0.54 as k ≥ 4, as shown in Fig. 4A. The minimum error for the top performing R^2^ models remains low, around 0.2 kcal/mol, with the average maximum error for the same models around 0.3 kcal/mol with the maximum error value observed for all systems falling below 1.0 kcal/mol at k=4 (Fig. S3). When we decompose the error between GP prediction and NetMHCIIpan prediction in Fig. 4B, certain residues have higher error, particularly residues Ala, Cys, Gly, and Pro. Meanwhile, residues Leu and Ser consistently have lower error and are predicted with higher accuracy than other residues. We further decomposed GP models by residue occurrence, calculating if certain residues occurred in top-scoring GP models more frequently than others, but we did not find large differences (Fig. S4). Similarly, scatterplot charts analogous to Fig. S2 do not reveal marked trends between R^2^, ΔG IQR, or residue position (Fig. S5). To evaluate whether GP regression can be generalized to perform binary classification of either enhanced or reduced binding affinity, we computed ROC curves and area under the curve (AUC) values, shown in Fig. 4C. Positive binding affinity values relative to the native antigen are considered reduced binding affinity while negative relative binding affinity values are considered enhanced. As shown, from the ROC curve of the raw data (Fig. 4C), GP regression has high accuracy for predicting the net binding affinity effects of mutations, with AUC values greater than 0.83 for all subsets with the maximum AUC of 0.96 for k=12. Other antigen binding prediction models including NetMHCIIpan 4.0, which use the entire antigen sequence, report AUC values ranging from 0.7 to 0.9 compared to experiment^12, 44-46^. Our method differs from these models by focusing on individual residue sites for guiding in-vivo or in-situ mutagenesis, incorporating predetermined information to increase the prediction accuracy in real time.

**Figure 4.**
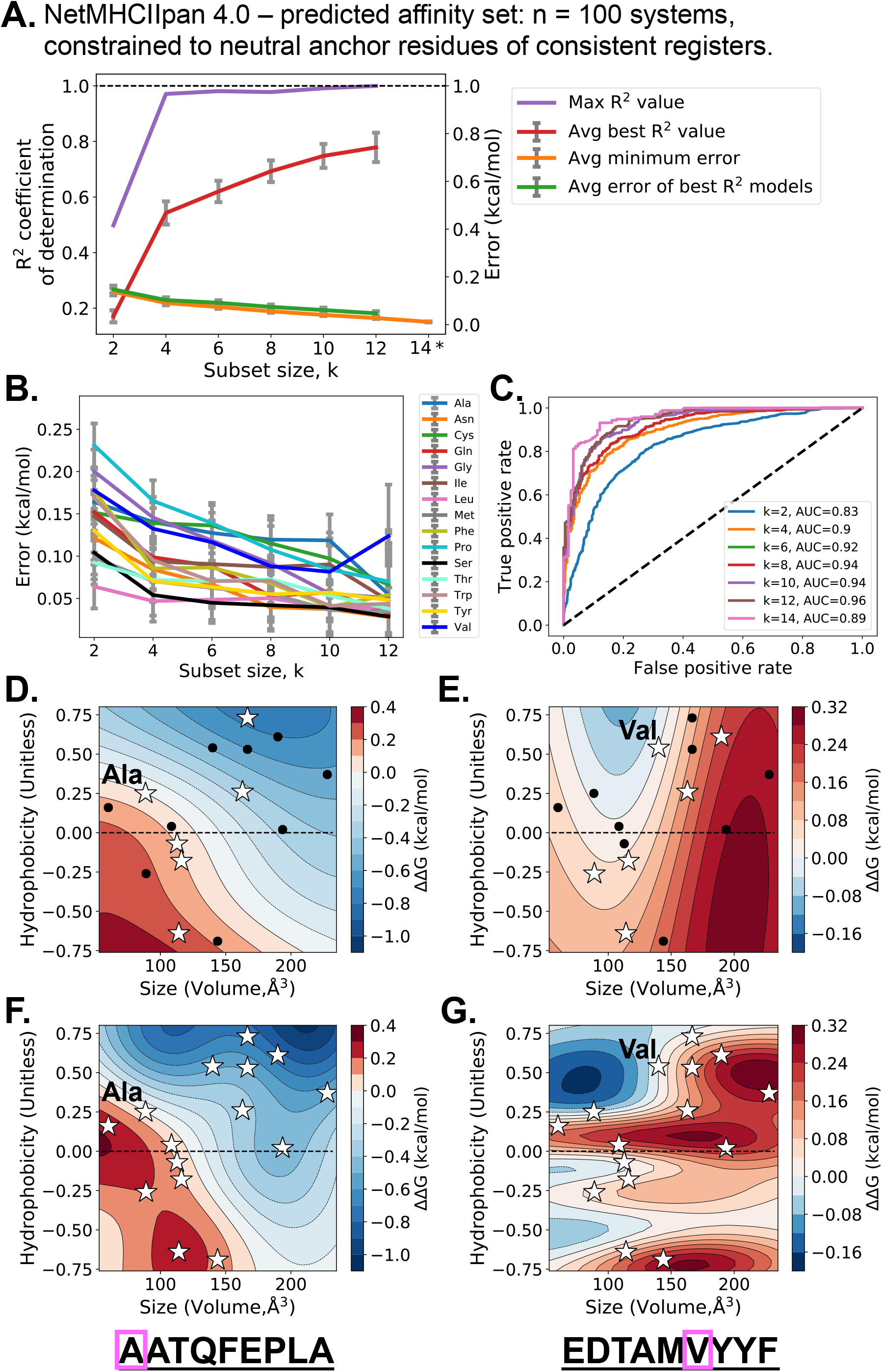
Gaussian process regression for register-constrained binding affinities from the NetMHCIIpan 4.0 server. **A**. Gaussian process regression R^2^ coefficient of determination and error for neutral residues at anchor residue positions. *k* is the subset size used for prediction, e.g., for k=4, 4 residue ΔG values were used to predict the remaining 11 residue ΔG values. Averages are taken across the n=100 systems. * For k=14 subset size, only 1 residue ΔG value is predicted, leaving R^2^ undefined. Error bars are 95% CI. **B**. Average error per residue for each k subset size for the top n=100 R^2^ models. Error bars are 95% CI. k=14 was excluded from B. as explained in A. **C**. ROC curve for multiple subset sizes, k with AUC values for classifying either enhanced or diminished binding affinity. **D-G**. Gaussian process regression binding affinity predicted surfaces for k=6 residue subsets (**D-E**.) and the full neutral residue sets (k=15, **F-G**.). **D**.**-E**. Surfaces represent maximum R^2^ models for k=6 residue subset shown as white stars with native residue labeled. Two examples are shown, (**D,F**.) predicting A1 mutants of **<u>AATQFEPLA</u>** binding to HLA-DQA10501-DQB10201 favoring larger, hydrophobic residues, and (**E,G**.) predicting V6 mutants of **<u>EDTAMVYYF</u>** binding to HLA-DQA10103-DQB10601 favoring smaller residues. Note that the full antigen sequence in D,F was included in the NETMHCIIpan training set, while the sequence in E,G was not.

Fig. 4D-G shows the GP-interpolated free energy surfaces for 2 systems, with both partial k=6 residue GP predictions (Fig. 4D-E), and complete set surfaces (Fig. 4F-G). Similar to GP-interpolated surfaces of the FEP values, the complete set surfaces preserve more detail than the partial subset surfaces, but the partial subsets are still able to capture broad trends and accurately predict mutant affinity values with 0.05 and 0.06 kcal/mol average error. Importantly, unlike other machine learning techniques which commonly favor large, hydrophobic residues for enhanced binding affinity, the GP prediction technique we employ here does not, as seen by the energy surfaces in Fig. 4E,G. Our GP prediction technique implicitly captures the actual MHCII-antigen binding interaction and then interpolates that interaction to other mutant residues. Hence, hydrophilic residues, and likewise more easily deliverable antigens, may also be favorably predicted.

## Discussion and future directions

GP regression can accelerate immunotherapy design by guiding mutagenesis and thereby decreasing the experimental and computational cost of determining MHCII binding affinity. We show that GP regression across a two-dimensional surface of neutral residue volume and hydrophobicity is sufficient to classify enhanced and reduced affinity mutations (AUC∼0.9+) and capture affinity trends across the mutational landscape (R^2^ coefficient of determination greater than 0.6 and low errors of 0.1-1.0 kcal/mol). This prediction method can be used concurrently with experimental and theoretical investigations to direct mutagenesis with real-time data. From a small residue subset, we can determine which residues are predicted to be favorable or have large uncertainty, and then select those particular residues to investigate next via experiment or computation. Furthermore, our GP prediction method benefits from two aspects. First, GP regression offers an estimate of error, so that we have knowledge of prediction confidence. Likewise, our GP regression method across a two-dimensional surface is intuitive; the impact of residue hydrophobicity and residue volume can be easily grasped rather than black box neural network methods or common QSAR techniques with large feature spaces.

For our GP prediction method, we implicitly assume antigen mutations do not affect binding register. Bound antigens are known to be dynamic^47-49^, and localized T-cell populations can recognize different register shifts of the same antigen^50^. However, for many ASI design studies, the target antigen, along with a probable binding register is known beforehand. Another assumption of our technique is that the bound antigens will not change binding conformations upon mutation, and/or that any residue size-dependent conformational changes will be captured by the GP model. For some proteins, single point mutations can drastically change structure^51-52^. But assuming the antigens are initially bound in the MHCII binding cleft, large structural changes are not anticipated, and if they do occur, we anticipate the binding effects will be implicitly captured by the GP-predicted free energy landscape.

We selectively chose to score GP regression models with coefficient of determination R^2^ values rather than Pearson product moment correlation coefficient R^2^ values. Indeed, as presented in Fig. S6, we tested scoring GP models by Pearson R^2^ values for the FEP systems of Fig. 2 and found quite high Pearson R^2^ values of 0.7 to 0.9 for subset size k=4. However, Pearson R^2^ values only quantify the existence of a linear relationship between the predicted affinities and the actual values. Pearson R^2^ values do not account for the variance of the data, so even though a GP model might have high Pearson R^2^ correlation, it may also have high error between the predicted affinities and actual values. The R^2^ coefficient of determination normalizes the error to the variance of the actual values, ensuring that high R^2^ coefficient of determination values also have low error between predicted affinities and actual values. Further, scikit-learn^53^, the machine learning library used to generate the GP models, implicitly scores models based on the R^2^ coefficient of determination values. We therefore conclude that the R^2^ coefficient of determination is a more logical metric of model accuracy and use it here for all investigations.

Our GP prediction method also opens doors for incorporating non-standard amino acids^54^ into antigen-specific immunotherapies. If a particular region of the interpolated GP landscape is predicted to be beneficial but not contain a standard amino acid, a non-standard amino acid in that region would be recommended for affinity testing. Extending ASI design to multiple mutation predictions simultaneously is also an area where GP regression-prediction methods might be useful. This would allow GP regression to gather information from alanine scan mutations^55^ and other multiple mutants with differential MHCII affinity.

## Methods

### Datasets

#### Free Energy Perturbation, FEP

FEP data were taken from references^2, 15^ based on mutagenesis calculations of antigens from a Type-1 Diabetes-implicated cell line. Briefly, two antigens were studied here, the X-idiotype: CARQE**<u>DTAMVYYFD</u>**YW and the healthy control: CARQ**<u>RFWSGPLFD</u>**YW. These antigens were modeled binding to HLA-DQ8, with the core epitope in bold underline. After 500ns of MD simulation to ensure stable pMHCII binding, FEP was conducted following protocols developed in previous works^41, 56^ using a custom short-range potential on NAMD2^57^. FEP was conducted over 34λ windows with at least 200ps/window and 5 replicas/mutation. All error bars are 95% CI unless otherwise noted.

#### Immune Epitope Database and Analysis Resource, IEDB

The IEDB webserver^42^ was accessed on Nov. 15, 2020 and the MHC-II binding dataset was downloaded from: http://tools.iedb.org/mhcii/download/. The IEDB dataset contains antigen binding data across multiple HLA types. An internal program was written to collect and collate antigens and their mutants, as well as convert binding affinity values to kcal/mol. Only antigens with binding affinity data for all 20 standard amino acids were selected for analysis. In total, 167 systems across 6 antigens and 11 HLA types met these criteria and were used for GP regression. The systems are presented in Table S1.

#### NetMHCIIpan 4.0 webserver

The NETMHCIIpan 4.0 webserver^12, 34, 58^ was accessed at http://www.cbs.dtu.dk/services/NetMHCIIpan/ on Feb. 1, 2021. Twenty antigens were selected for prediction based on their occurrence in the IEDB as well as implication in Type-1 Diabetes. The systems are presented in Table S2. The epitopes were register-constrained during the NetMHCIIpan prediction by truncating the flanking domains, leaving only the predicted 9-residue epitope core. Only mutants for anchor residues 1, 4, 6 and 9 were computed, leading to a total of 20 antigens x 4 anchors = 100 systems. The binding affinity rather than the eluted ligand metric was used for affinity values.

### Gaussian Process (GP) Regression

GP models were generated using scikit-learn’s machine learning python library^53^. A constant kernel combined with an RBF kernel was used for model generation. Median kernel parameters are presented in Fig. S7. For each system, all possible residue subset combinations were tested for GP generation, only saving the top 100-best scoring and minimum error models.

## Acknowledgements

The authors would like to thank Leili Zhang, Guojing Cong, Giacomo Domeniconi, Chih-Chieh Yang, Ruhong Zhou, Jeffrey K Weber, and Sangyun Lee for insightful discussions. SHC would like to acknowledge Program Development funding from Oak Ridge National Laboratory which helped spur this collaboration.

This project has been funded in whole or in part with Federal funds from the National Cancer Institute, National Institutes of Health, under Contract No. HHSN261200800001E. The content of this publication does not necessarily reflect the views or policies of the Department of Health and Human Services, nor does mention of trade names, commercial products, or organizations imply endorsement by the U.S. Government.

## Author Contributions

DRB and SHC designed and conceived the study as well as ran experiments. DRB performed data analysis and generated the figures. DRB and SHC wrote the manuscript.

## Supplementary Information

**Figure S1.**
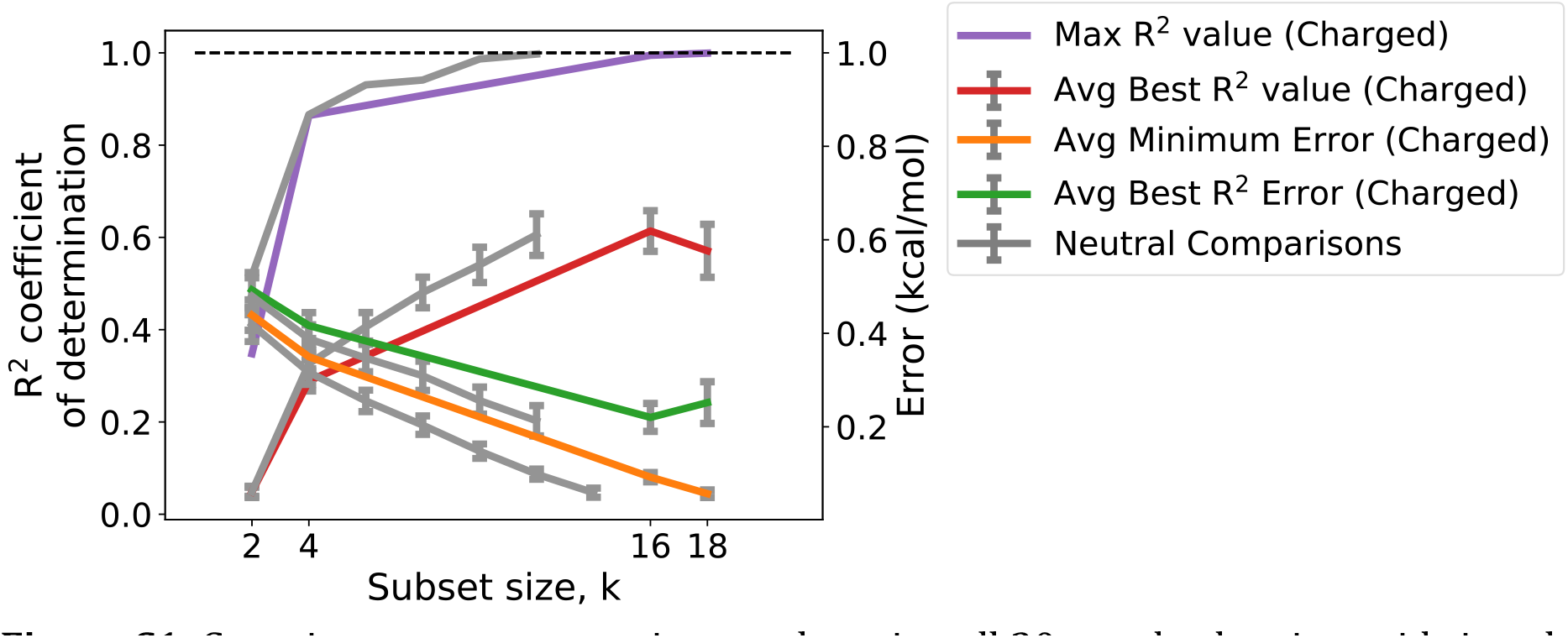
Gaussian process regression results using all 20 standard amino acids in color, compared to using 15 neutral amino acids in gray. Data is experimentally determined binding affinities from the Immune Epitope Database (IEDB). R^2^ is the coefficient of determination. For the 20 standard amino acids, GP regression was evaluated for k=2,4,16, and 18 subset sizes, while neutral residues were evaluated at k=2,4,6,8,10,12, and 14. Averages are taken across the n=167 systems. The neutral comparison plots in gray are the same as presented in Figure 3 in main text. Error bars are 95% confidence intervals.

**Figure S2.**
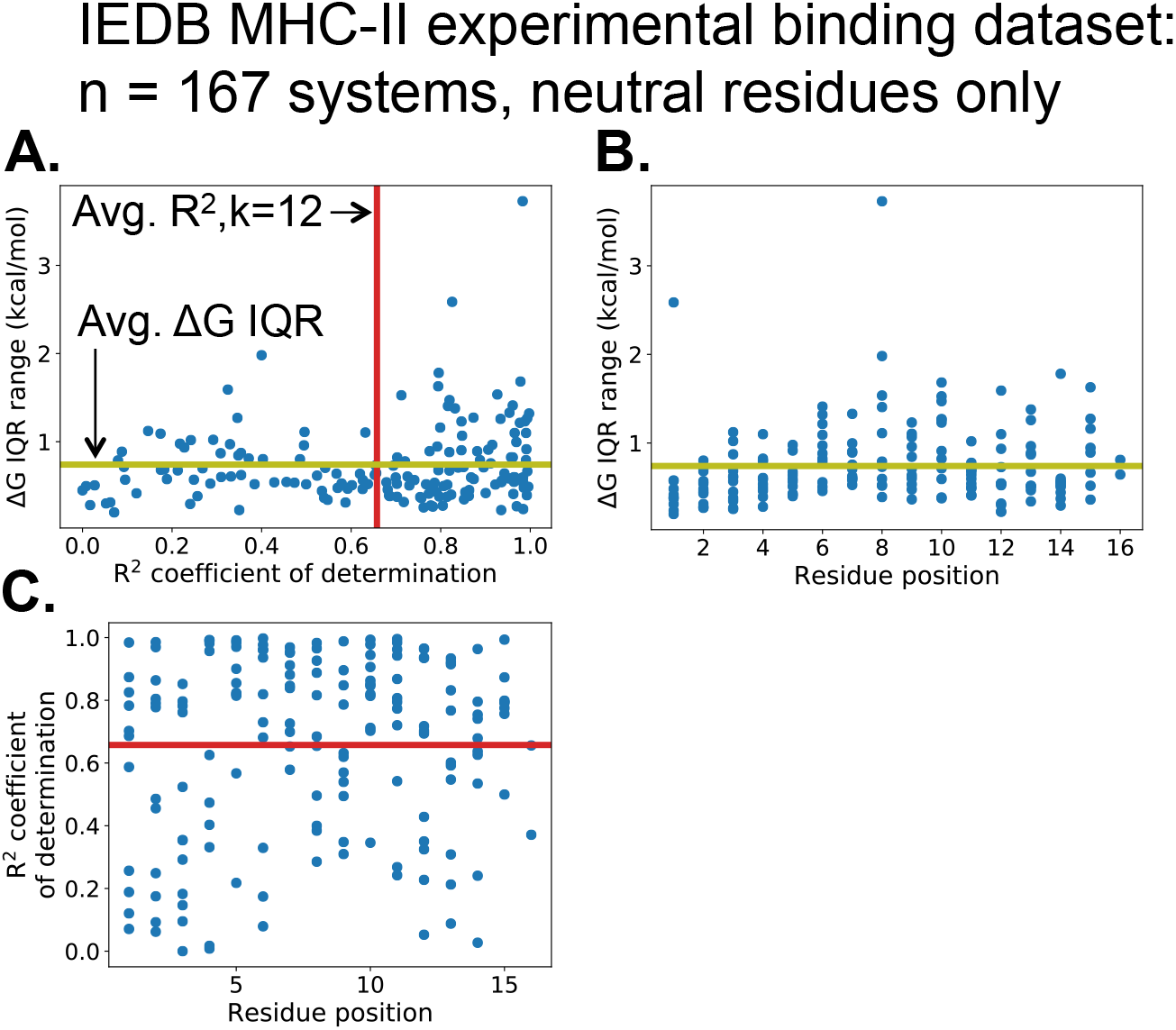
ΔG interquartile range (IQR) and Gaussian process regression performance for the n=167 experimentally determined systems from the Immune Epitope Database (IEDB). **A**. Binding affinity ΔG IQR for n=167 systems as a function of Gaussian process regression R^2^ coefficient of determination values for the k=12 subset size. **B**. Binding affinity ΔG IQR for n=167 systems as a function of residue position in the MHCII antigen. **C**. Gaussian process regression R^2^ values for the k=12 subset size across residue position in the MHCII antigen. **A-C**. Avg ΔG IQR from expt. and Avg R^2^ values for k=12 subset size is shown in olive and red lines respectively.

**Figure S3.**
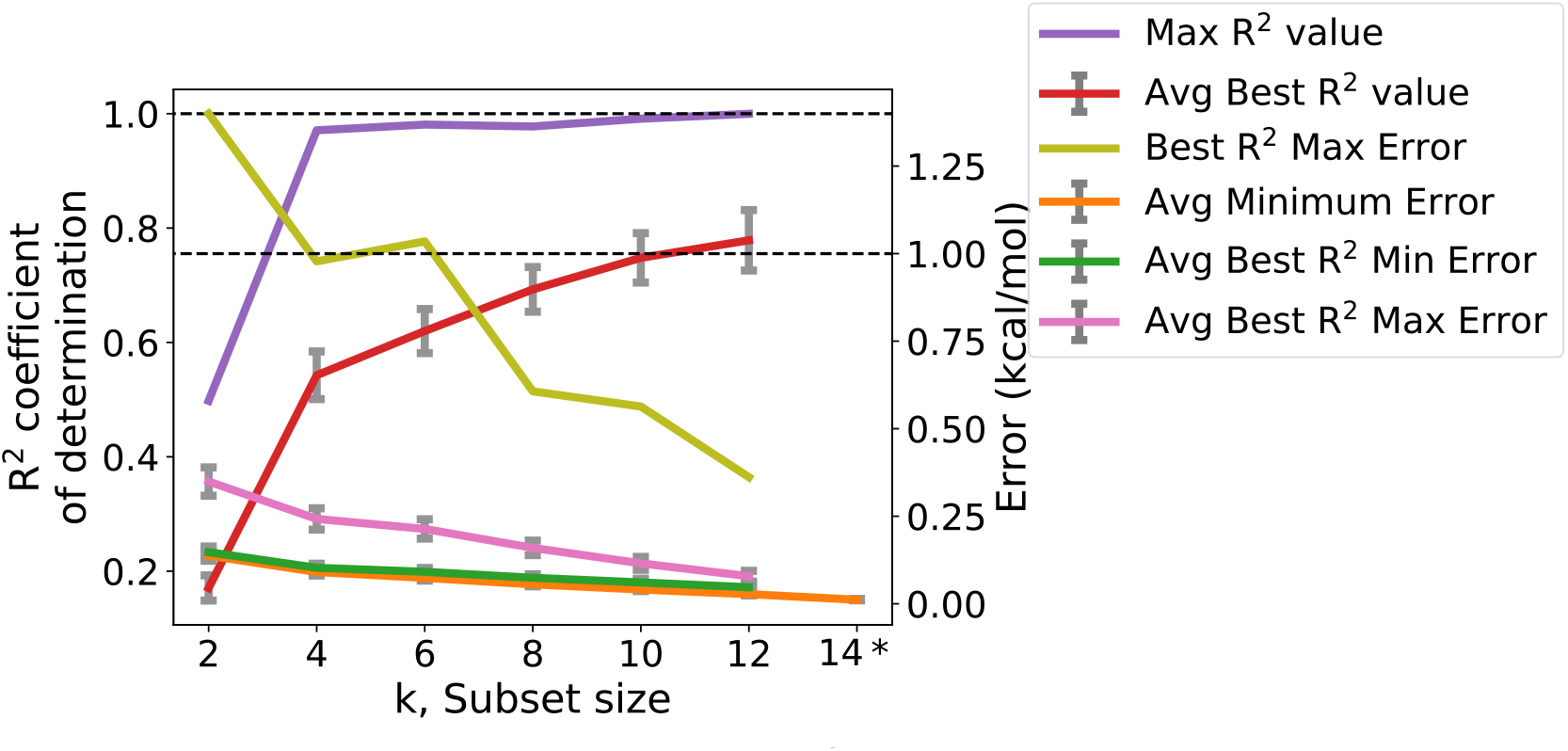
Gaussian process regression R^2^ coefficient of determination and error for only neutral residues (15 total neutral residues) using the NETMHCIIpan dataset. k is the subset size used for prediction, e.g. for k=4, 4 residue ΔG values were used to predict the remaining 11 residue ΔG values. Averages are taken across the n=100 register-constrained systems predicted from NetMHCIIpan 4.0. Error bars are 95% CI. “Best R^2^ Max Error” means the highest residue error of the top scoring models for the n=100 systems at k subset size.

**Figure S4.**
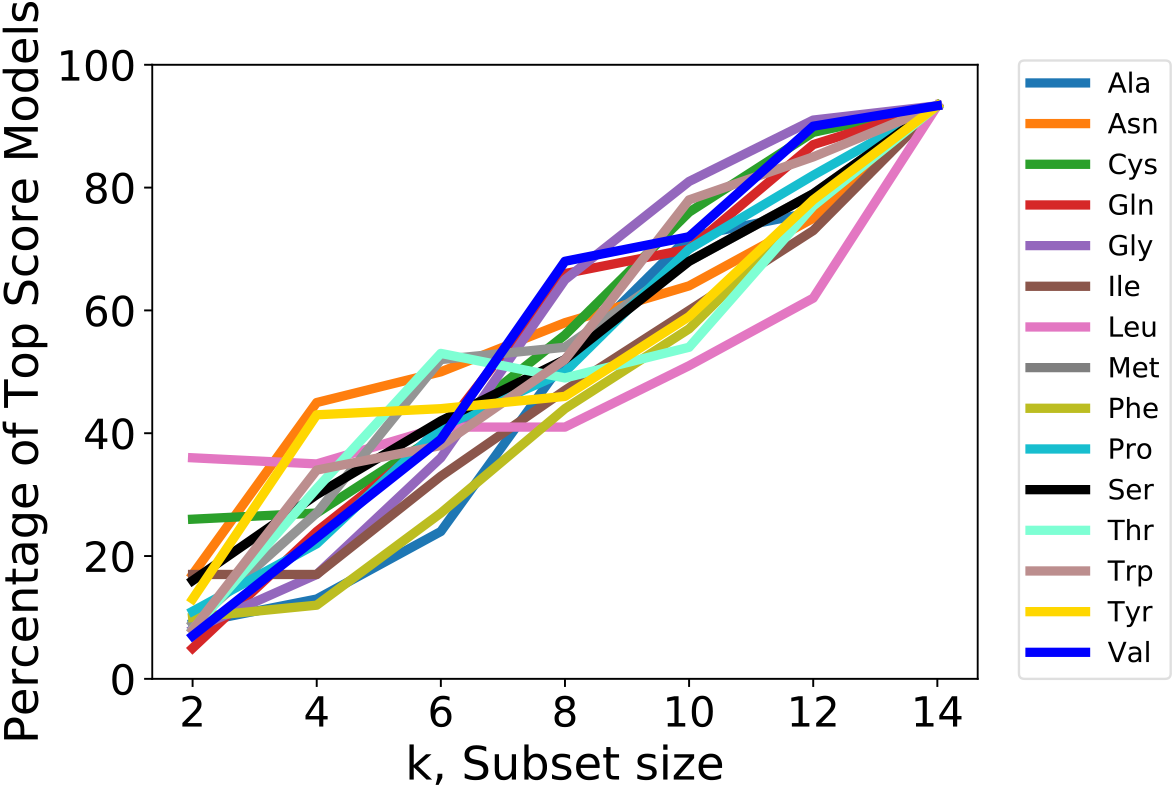
Residue occurrence in Gaussian process regression models across subset size, k for the NetMHCIIpan dataset. Note: only neutral residues are shown, charged residues were excluded from our analysis.

**Figure S5.**
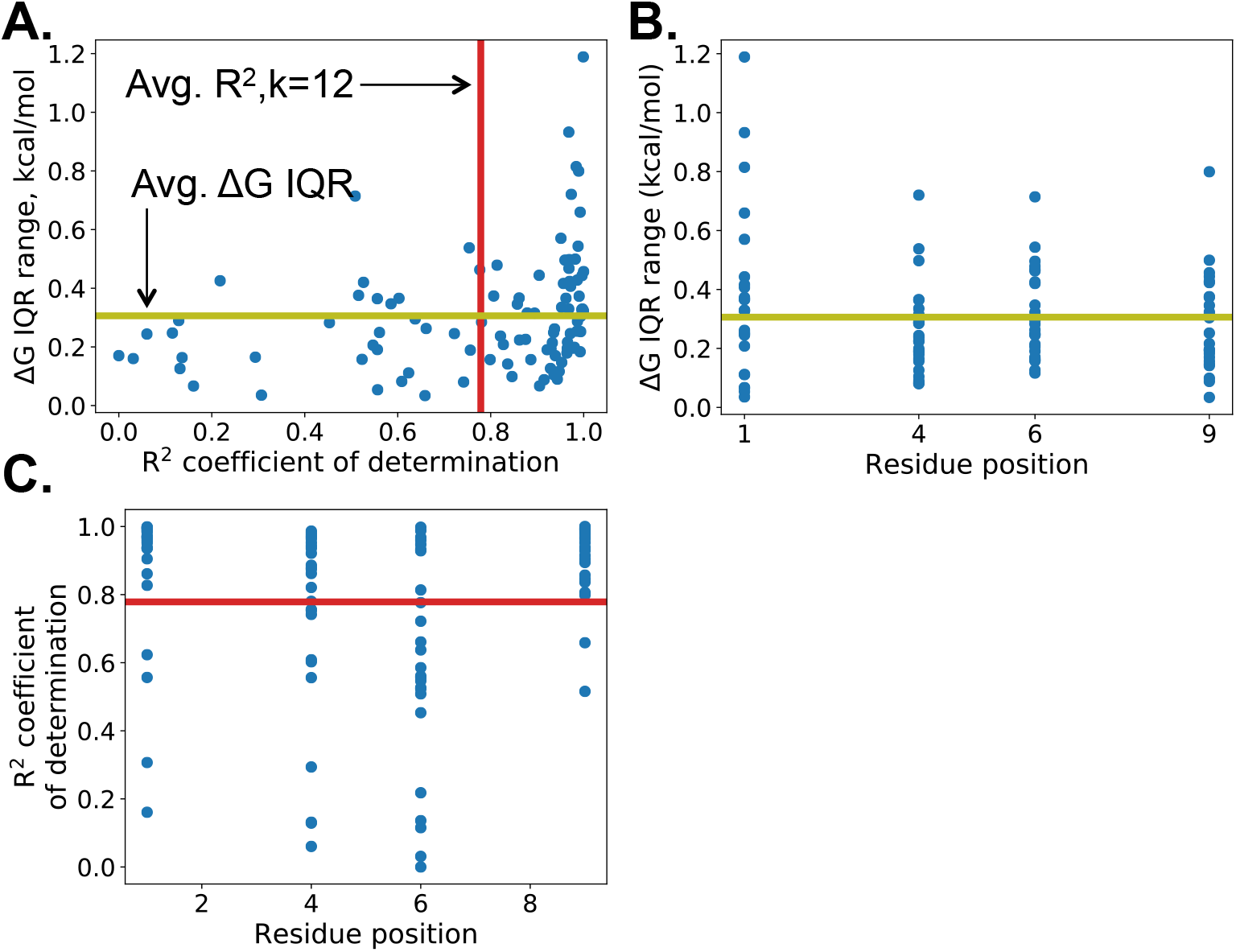
ΔG interquartile range (IQR) and Gaussian process regression performance for the n=100 register-constrained NetMHCIIpan 4.0-predicted set. **A**. Binding affinity ΔG IQR for n=100 systems as a function of Gaussian process regression R^2^ values for the k=12 subset size. R^2^ is the coefficient of determination. **B**. Binding affinity ΔG IQR for n=100 systems as a function of residue position in the MHCII antigen. **C**. Gaussian process regression R^2^ values for the k=12 subset size across residue position in the MHCII antigen. **A-C**. Avg ΔG IQR from expt. and Avg R^2^ values for k=12 subset size is shown in olive and red lines respectively.

**Figure S6.**
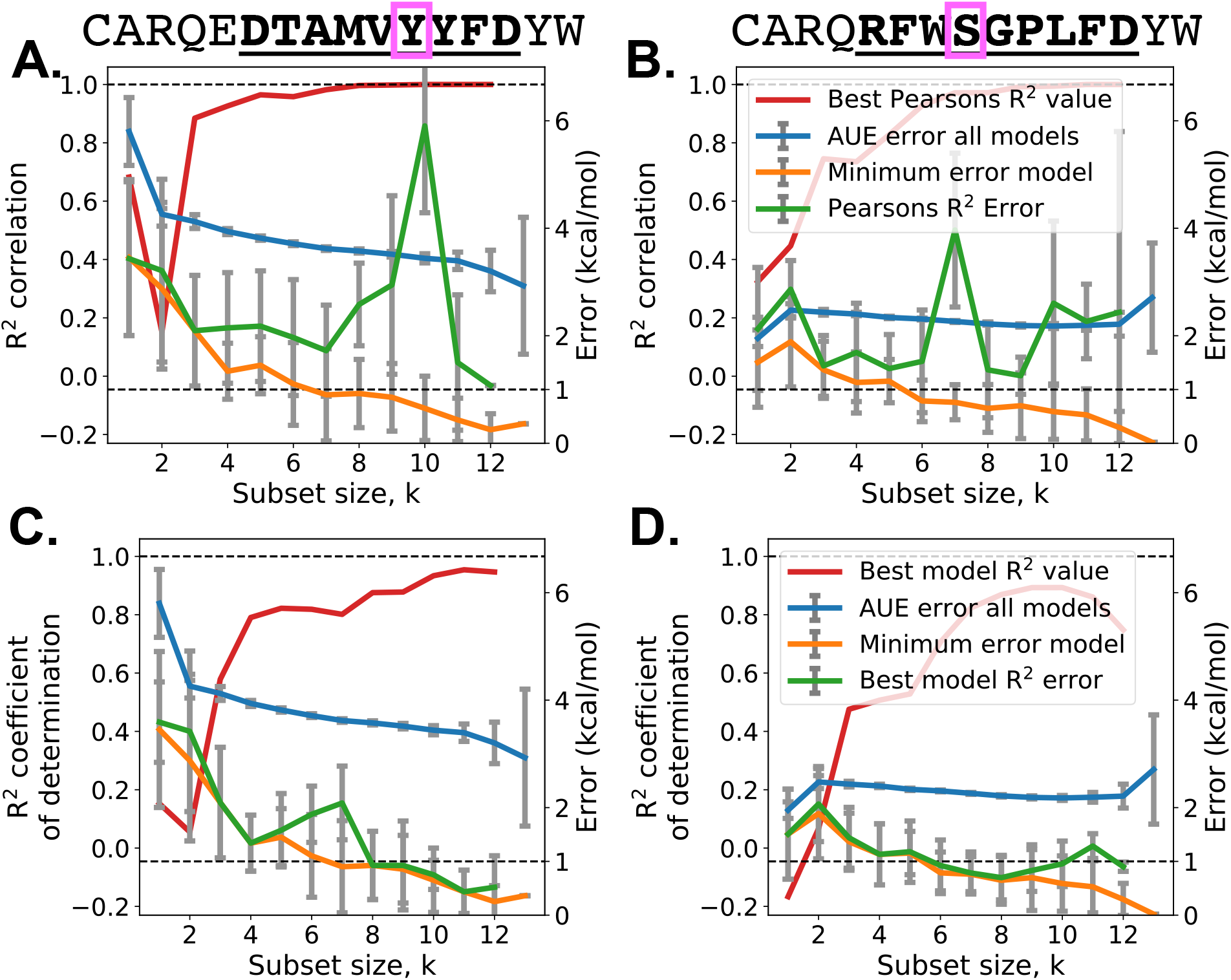
Gaussian process results comparing scoring by Pearson’s product moment correlation coefficient (**A-B**) and the coefficient of determination (**C-D**). Note that systems are the same as shown in Fig. 2. Error bars represent 95% confidence intervals. Although the Pearson correlation coefficient is high, the variance of the data is not accounted for, hence the overall error does not decrease with increasing correlation coefficient values. In contrast, the R^2^ coefficient of determination does account for data variance or spread, and thus higher coefficient of determination values generally results in lower errors.

**Figure S7.**
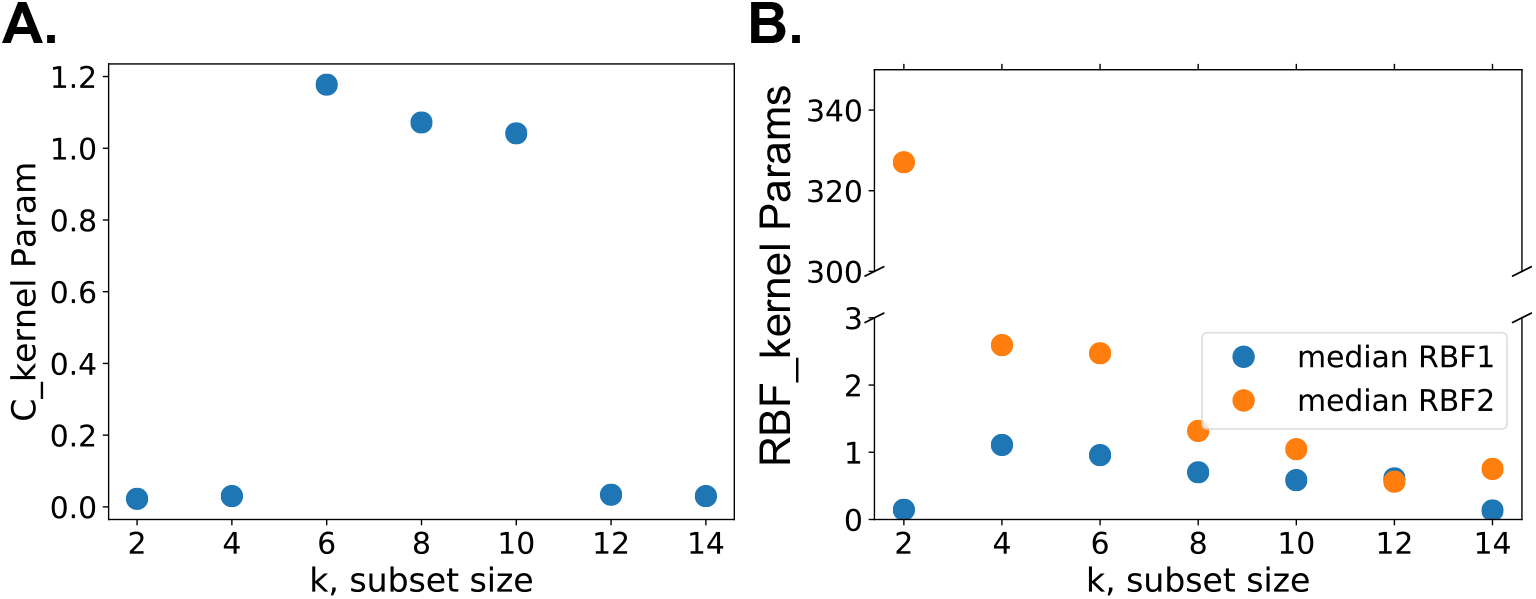
Gaussian process regression kernel parameters found from the NetMHCIIpan dataset. A constant kernel and a radial basis function kernel were used from scikit-learn’s framework. **(A)** The C kernel parameter median value and **(B)** the RBF kernel parameters median values across k, subset size: RBF1=RBF parameter volume, RBF2=RBF parameter hydrophobicity.

**Table S1.**
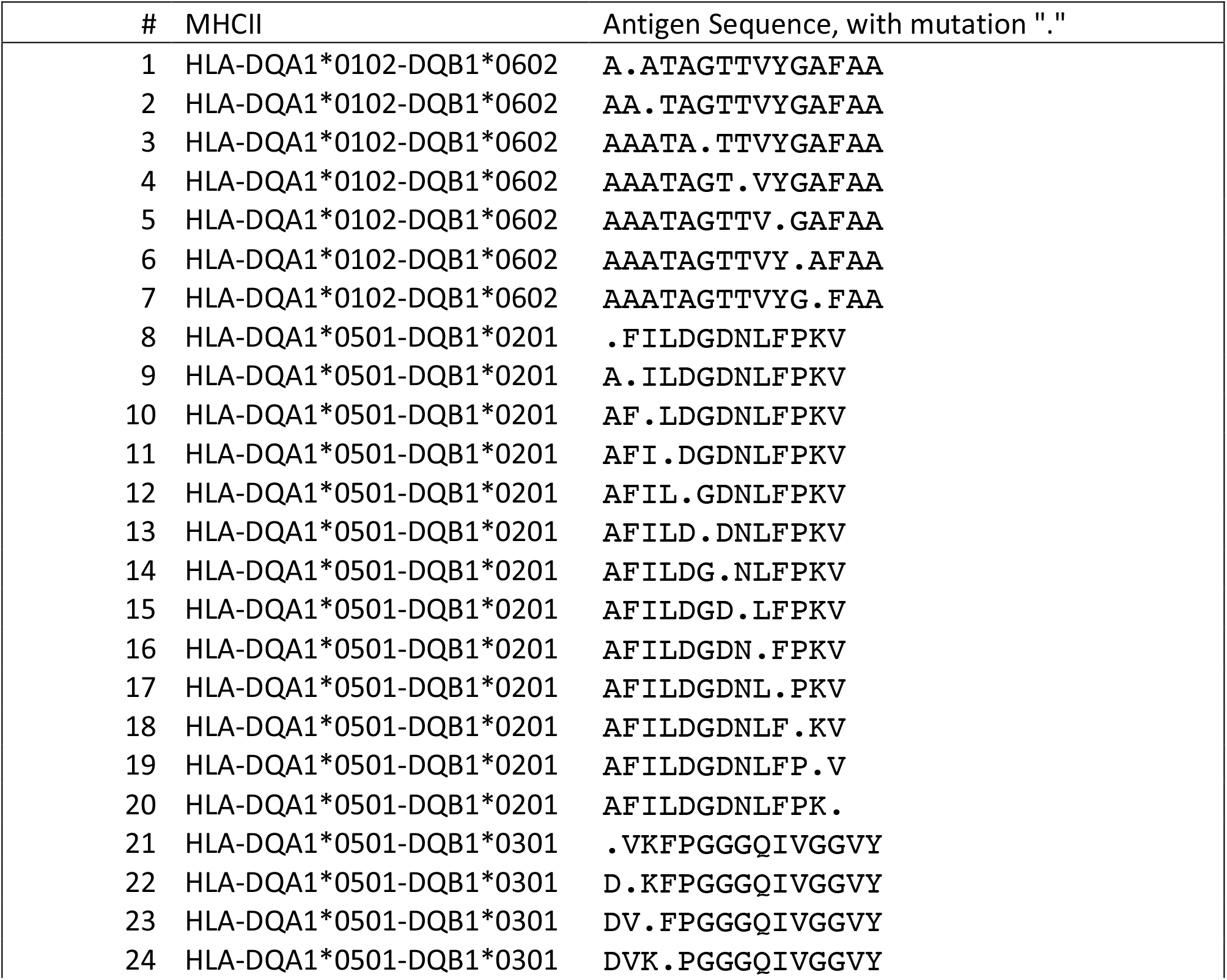
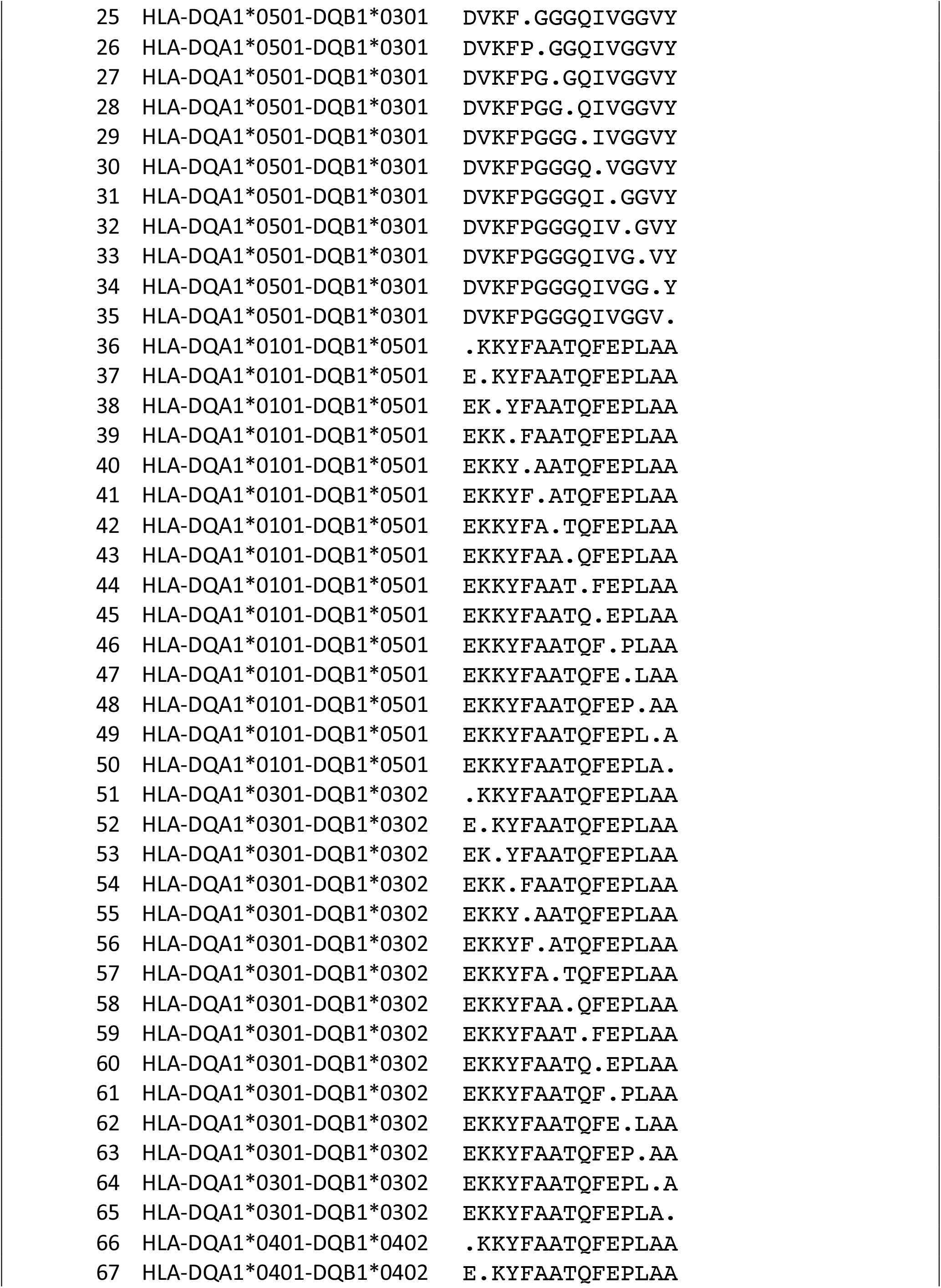
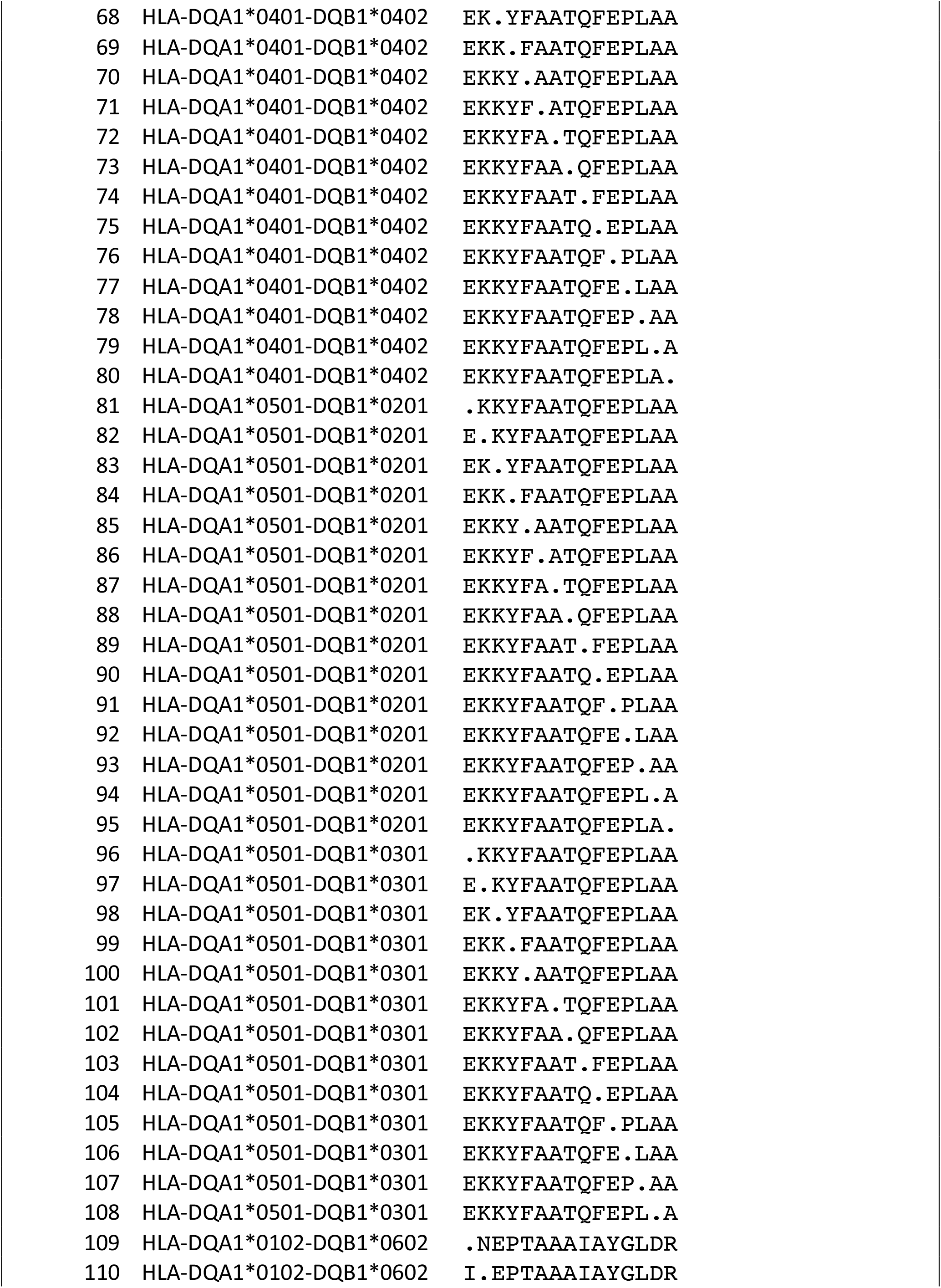
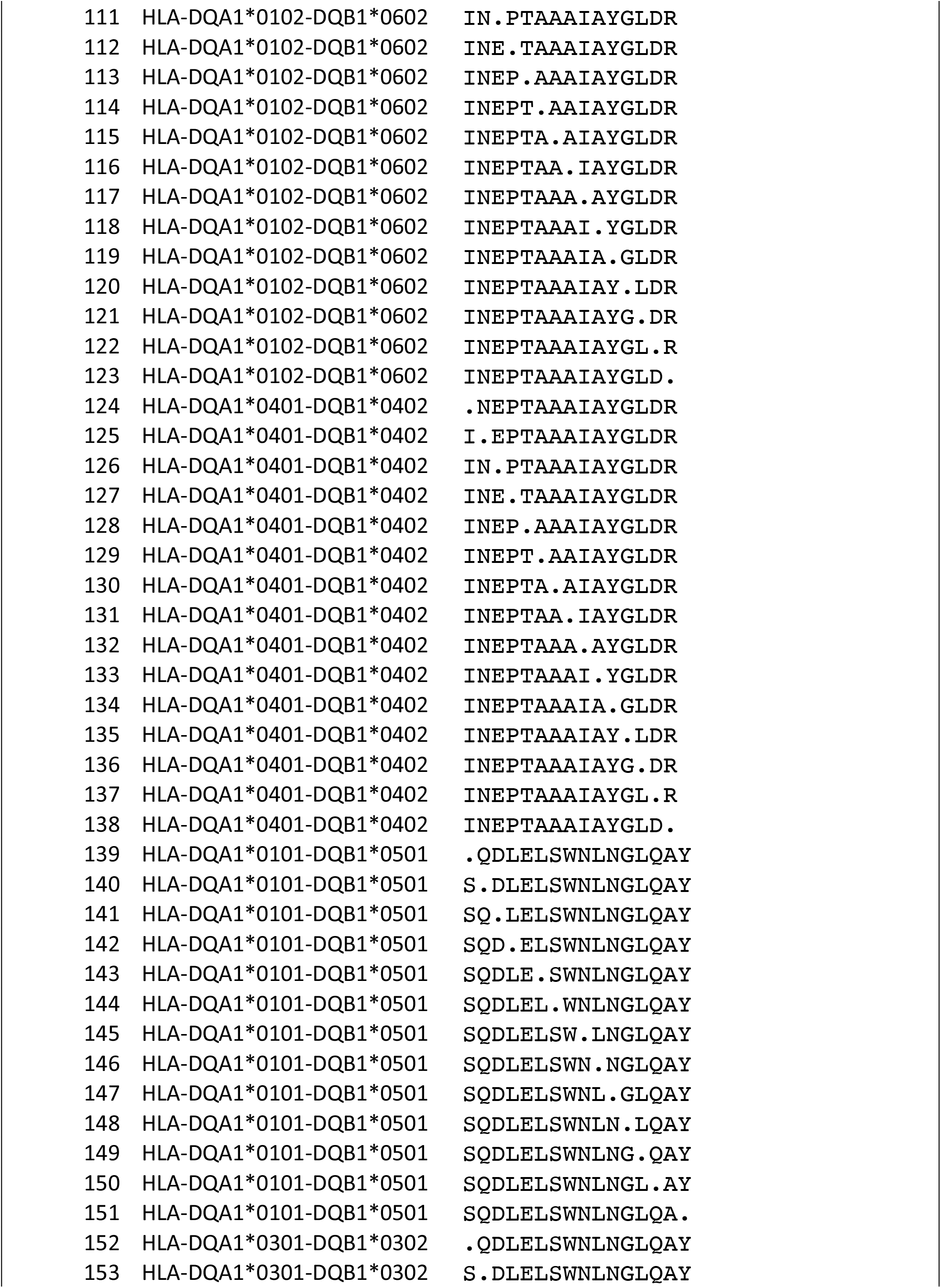
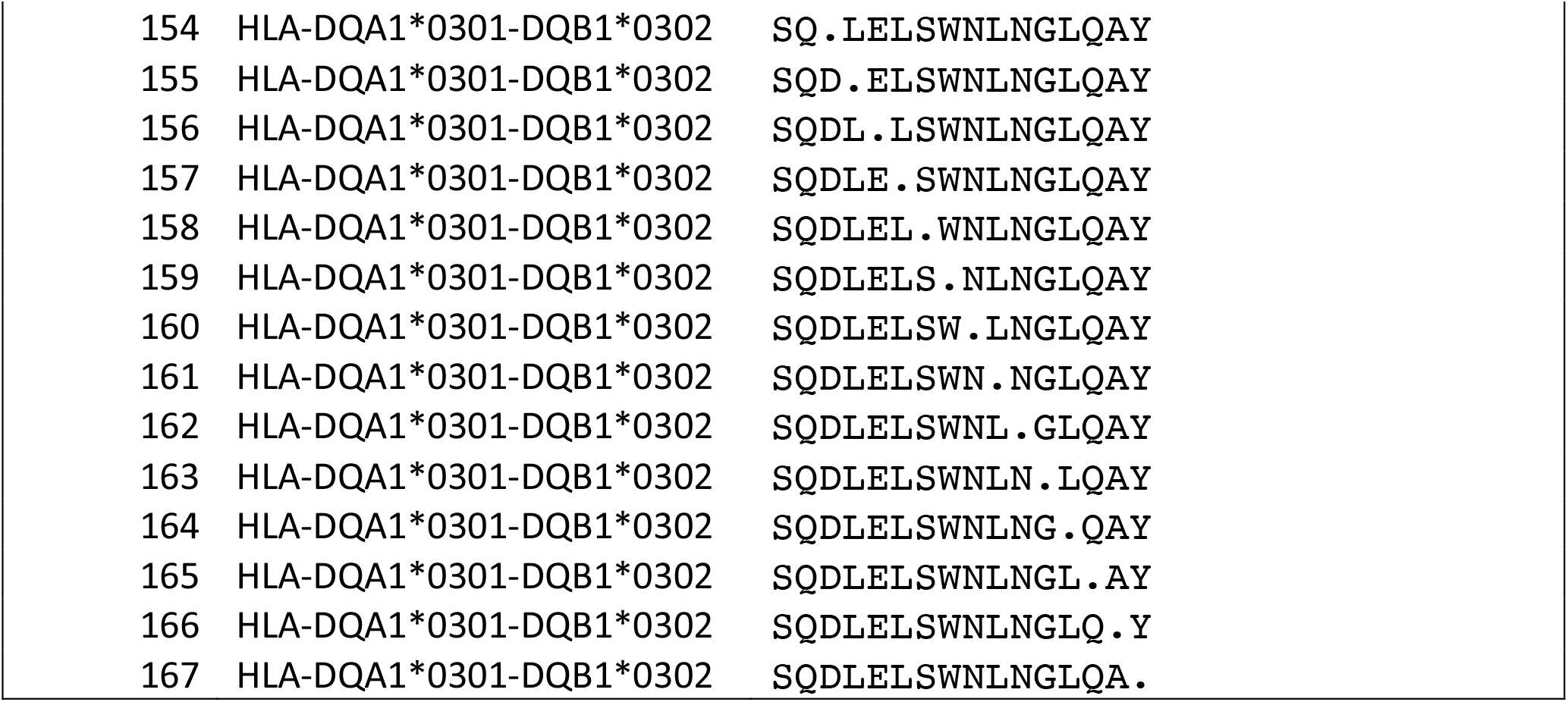
IEDB dataset

**Table S2.**
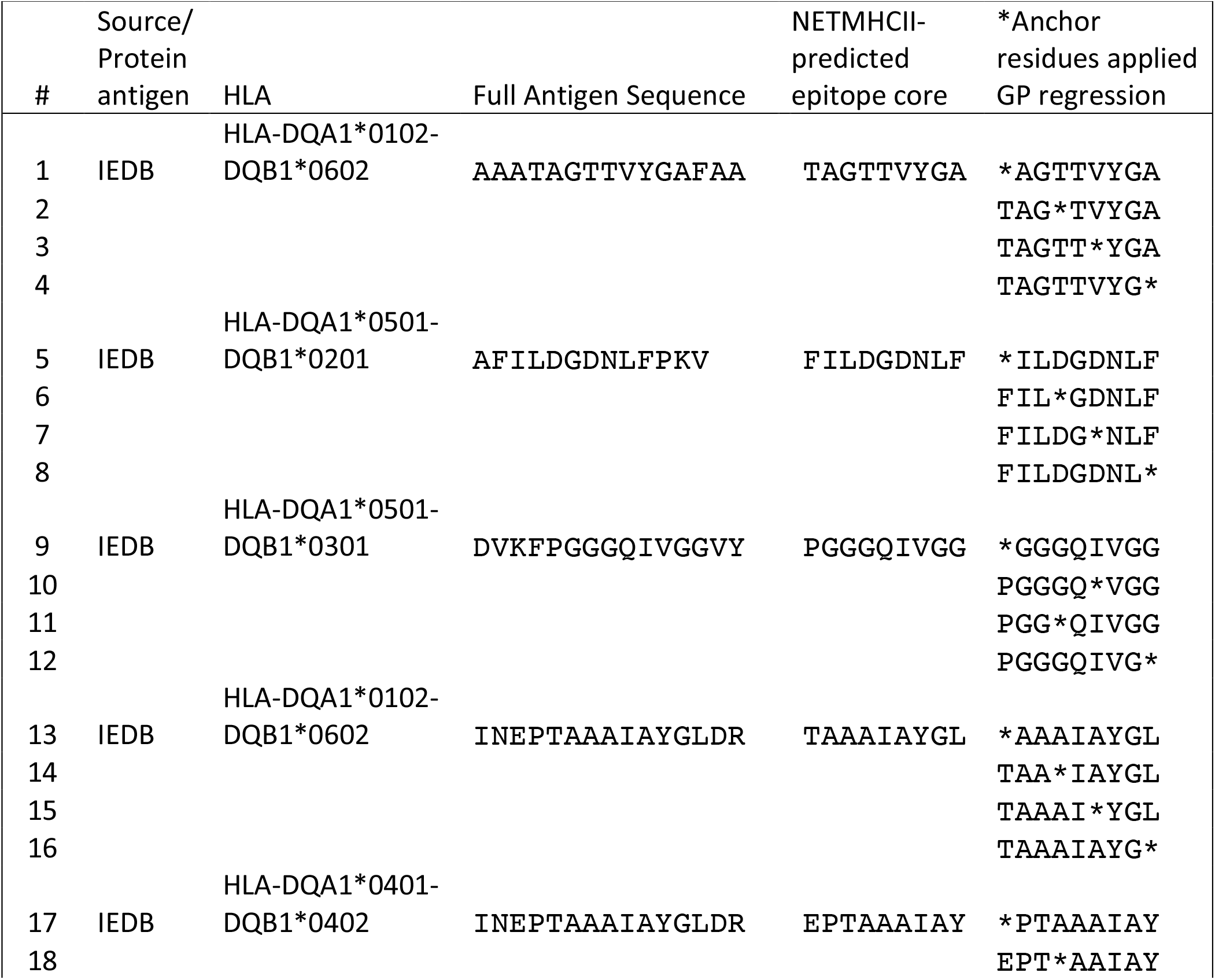
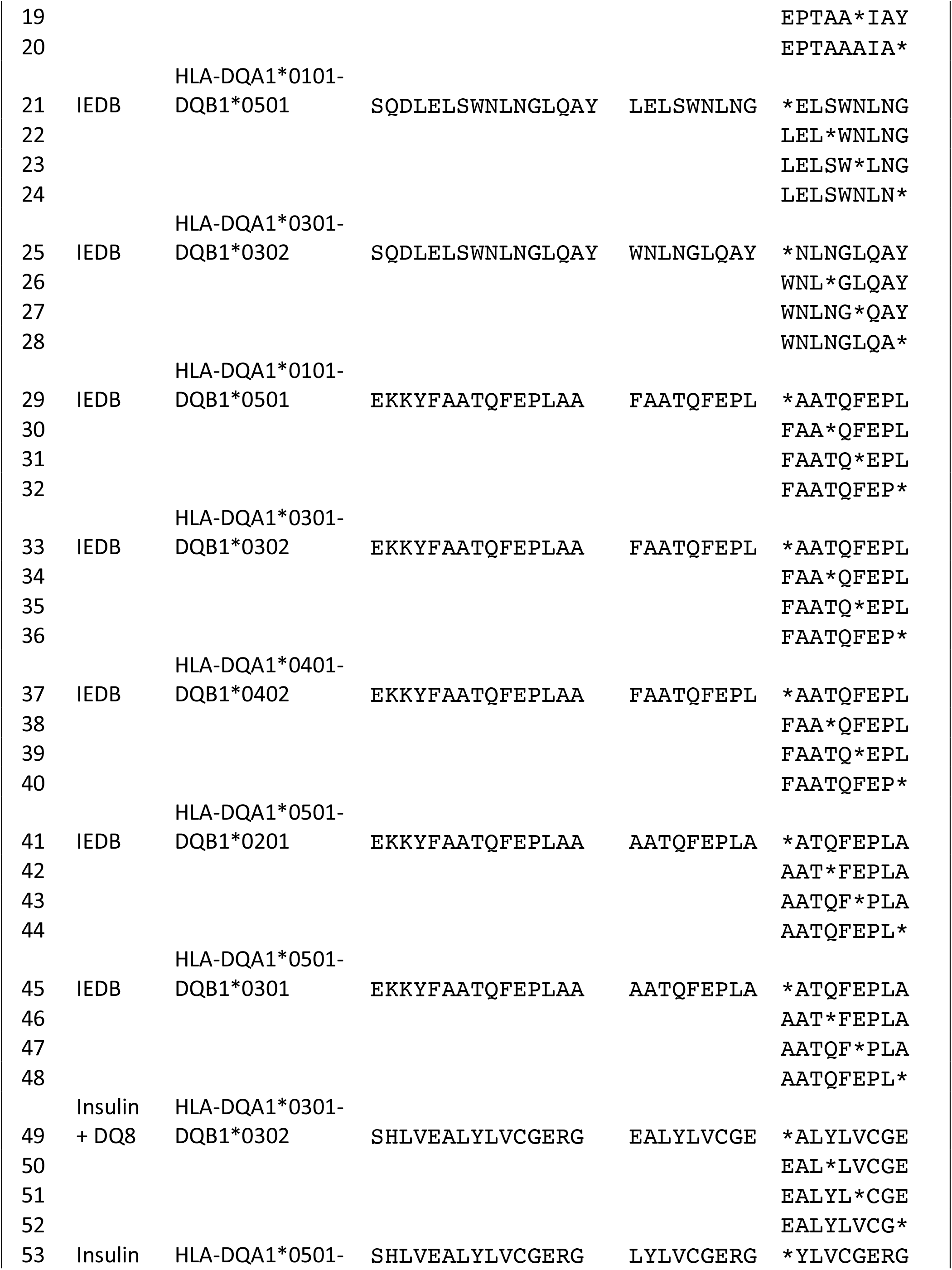
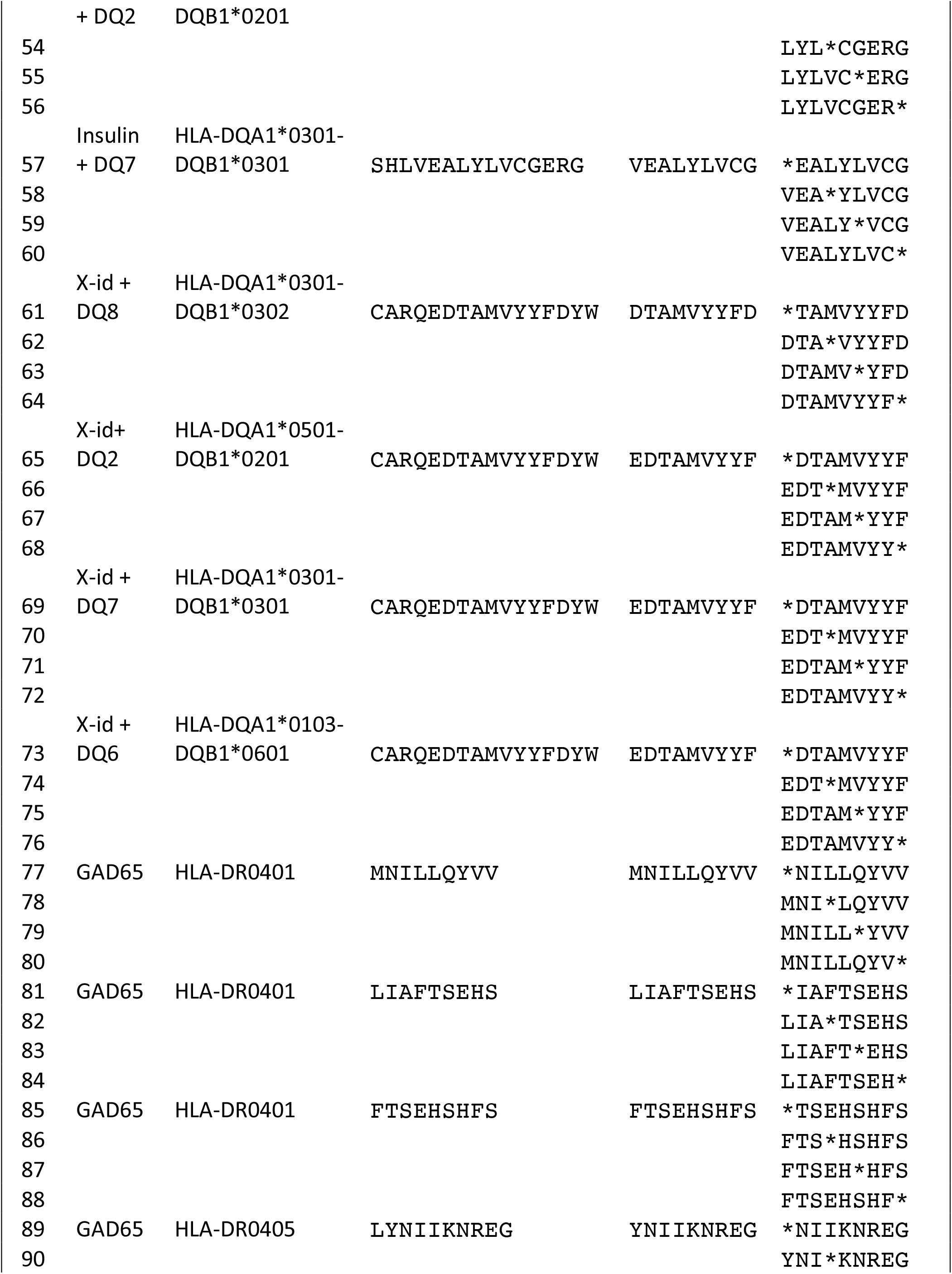
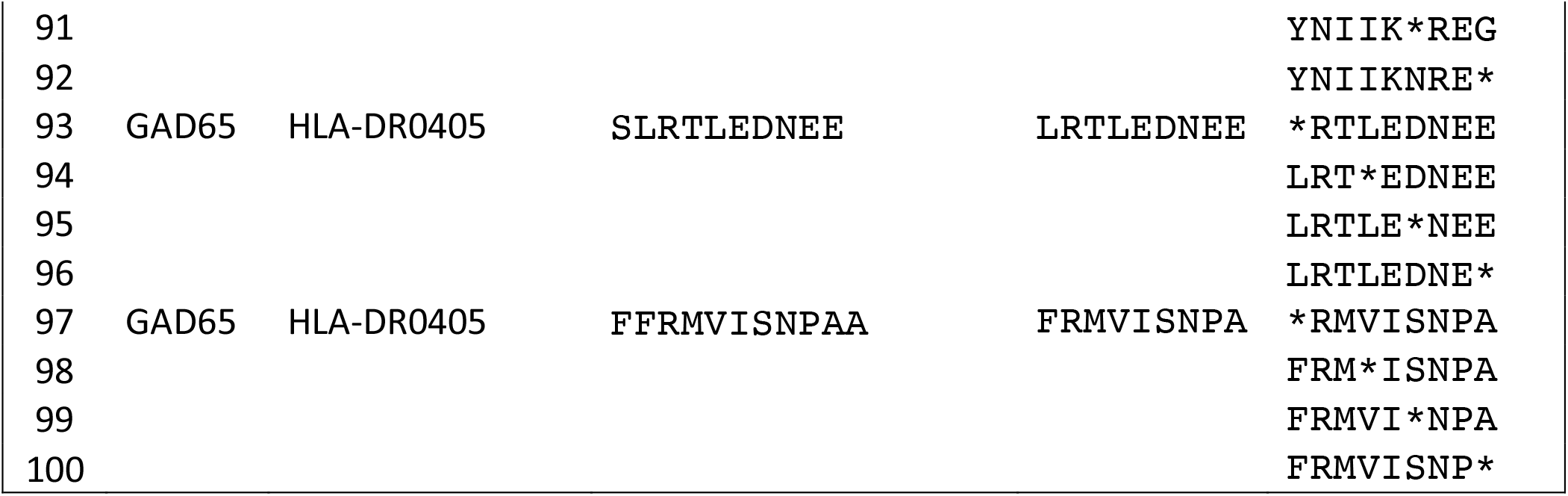
NetMHCIIpan 4.0 sequence dataset.

## Notes

### Competing Interest Statement

The authors have declared no competing interest.

